# LEMONmethyl-seq: Targeted long-read DNA methylation profiling reveals dynamics of CRISPR epigenome editing and endogenous DNA methylation patterns

**DOI:** 10.64898/2026.02.24.707761

**Authors:** Anna E. Christenson, Nikita S. Divekar, Justin P. Lubin, Luis G. Palma, Peter J. Colias, Rithu K. Pattali, Da Xu, Akane Hubbard, Katie Lin, Ngan T. Phan, Bernardo D. Moreno, Sarah E. Chasins, S. John Liu, James K. Nuñez

## Abstract

**BACKGROUND:** DNA methylation is the most prevalent epigenetic modification in human cells and undergoes dynamic changes during cell differentiation, disease progression, and aging. Here, we introduce <u>L</u>ocus-<u>E</u>nriched <u>M</u>apping <u>O</u>f <u>N</u>ucleotide methylation (LEMONmethyl-seq): an optimized, cost-effective pipeline for single-nucleotide detection of DNA methylation using locus-specific amplification and long-read DNA sequencing.

**RESULTS:** We apply LEMONmethyl-seq to profile DNA methylation of endogenous gene promoters across different cell types along with DNA methylation establishment and long-range propagation induced by CRISPR epigenome editing technologies. We profile dynamic changes in DNA methylation patterns on transposable element genomic loci during global epigenetic resetting in stem cells, and we identify site-specific enrichment of non-canonical CpH methylation on genomic sites in stem cells and cultured neurons. Lastly, we apply LEMONmethyl-seq to profile DNA methylation across the *MGMT* promoter, a clinical biomarker for glioblastoma. We identify additional differentially methylated sites correlated with chemotherapeutic sensitivity, which may be clinically relevant.

**CONCLUSIONS:** Together, LEMONmethyl-seq serves as a cost-effective, long-read DNA methylation sequencing pipeline that advances methods for detecting DNA methylation patterns and dynamics in mammalian cells. We envision its broad use for studying chromatin pathways, diagnostics, and therapeutic applications.

## BACKGROUND

DNA methylation is the most abundant chemical modification on the human genome, typically defined as methylation at the 5’ position of cytosines [1,2]. DNA methylation patterns undergo dynamic changes during early embryonic development and cell differentiation that contribute to gene expression patterns, genome organization, and cell type identity [2–5]. Aberrant DNA methylation patterning is a hallmark of cancer and many neurodevelopmental diseases, such as Rett Syndrome and Tatton-Brown-Rahman Syndrome [6–12]. Recently, DNA methylation has emerged as a biomarker for biological aging by measuring DNA methylation changes at specified sites on the human genome [13–20]. Thus, methods for measuring DNA methylation patterns at defined genomic sites have broad applications for fundamental research, diagnostics, and therapeutic applications.

The most common methods to profile site-specific DNA methylation include bisulfite conversion, enzymatic methyl (EM) conversion, and direct readout of endogenous methylation with third-generation sequencing technologies [21–24]. Bisulfite conversion utilizes sodium bisulfite to deaminate unmethylated cytosines to uracil, and the resulting bisulfite-converted DNA is PCR-amplified to enrich for a region of interest [21]. During PCR amplification, uracil is replaced with thymine, allowing cytosine positions in the original sample to be classified as methylated (cytosine) or unmethylated (thymine). Conventionally, the amplified PCR product is ligated into a plasmid vector, bacteria are transformed, individual colonies are picked the next day, and plasmid DNA is extracted for Sanger sequencing of the ligated product. Typical DNA methylation data from bisulfite-PCR is obtained from 5 to 15 Sanger reads. Second-generation high-throughput sequencing is an alternative approach to improve read depth across the PCR amplicon and increase confidence in DNA methylation quantification [25,26] but is bottlenecked by high sequencing costs. Additionally, bisulfite-PCR is optimal for amplifying 200-300 base pair (bp) fragments due to the high levels of DNA fragmentation induced by sodium bisulfite treatment, thus limiting its use for DNA methylation patterns across longer genomic segments [27].

To reduce DNA damage allowing for generation of kilobase amplicons, EM conversion for DNA methylation detection was developed recently [22]. First, hydroxymethyl (5hmC) and methyl (5mC) cytosines are oxidized by the TET2 enzyme into 5-carboxycytosine (5caC) or 5-glucosylmethylcytosine (5gmC). These chemical modifications protect the cytosines from later deamination by the APOBEC enzyme, which converts unmodified cytosine into uracil. Similar to bisulfite conversion, EM-converted DNA can be PCR-amplified for target site enrichment and sequenced to detect methylated and unmethylated cytosines. While prior work has shown that EM-seq coupled with PCR enrichment and second- or third-generation sequencing is feasible [28–30], the approach has not been widely adopted due to limited proof-of-concept across experimental applications, inconsistent amplification across experiments, and computational hurdles.

Third-generation sequencing methods, such as Oxford Nanopore and PacBio sequencing, allow for long-read sequencing and endogenous DNA methylation calling [31–38]. PCR amplification results in loss of DNA methylation information, requiring alternative methods for target-site enrichment. Adaptive sampling is one such method for Oxford Nanopore sequencing, though it is computationally intensive and can lead to pore degradation [39,40]. Additionally, recent studies have noted false positive methylation calling, especially for noncanonical CpH sites (CpC, CpA, CpT) [41–43]. Although methylation at CpH sites is rare in mammalian cells, CpH methylation is a prominent and dynamic epigenetic mark in mammalian neurons and stem cells, where it regulates gene expression and is closely linked to neurodevelopmental processes and pluripotent identity [44–49].

We sought to develop a reproducible, streamlined, and cost-effective pipeline to profile DNA methylation at target sites with long-read DNA sequencing. Here, we introduce <u>L</u>ocus-<u>E</u>nriched <u>M</u>apping <u>O</u>f <u>N</u>ucleotide methylation (LEMONmethyl-seq), an optimized experimental and computational pipeline for locus-specific, single-nucleotide detection of DNA methylation across thousands of base pairs in a single sequencing run (**Fig. 1a**). We apply LEMONmethyl-seq to profile DNA methylation across various experimental contexts, including following CRISPR-based DNA methylation editing, across transposable elements during stem cell naïve resetting, and at a clinically relevant biomarker, highlighting its many applications and robust performance (**Fig. 1b**). LEMONmethyl-seq serves as a cost-effective, long-read DNA sequencing pipeline that advances methods for detecting DNA methylation patterns and dynamics in mammalian cells.

**Figure 1.**
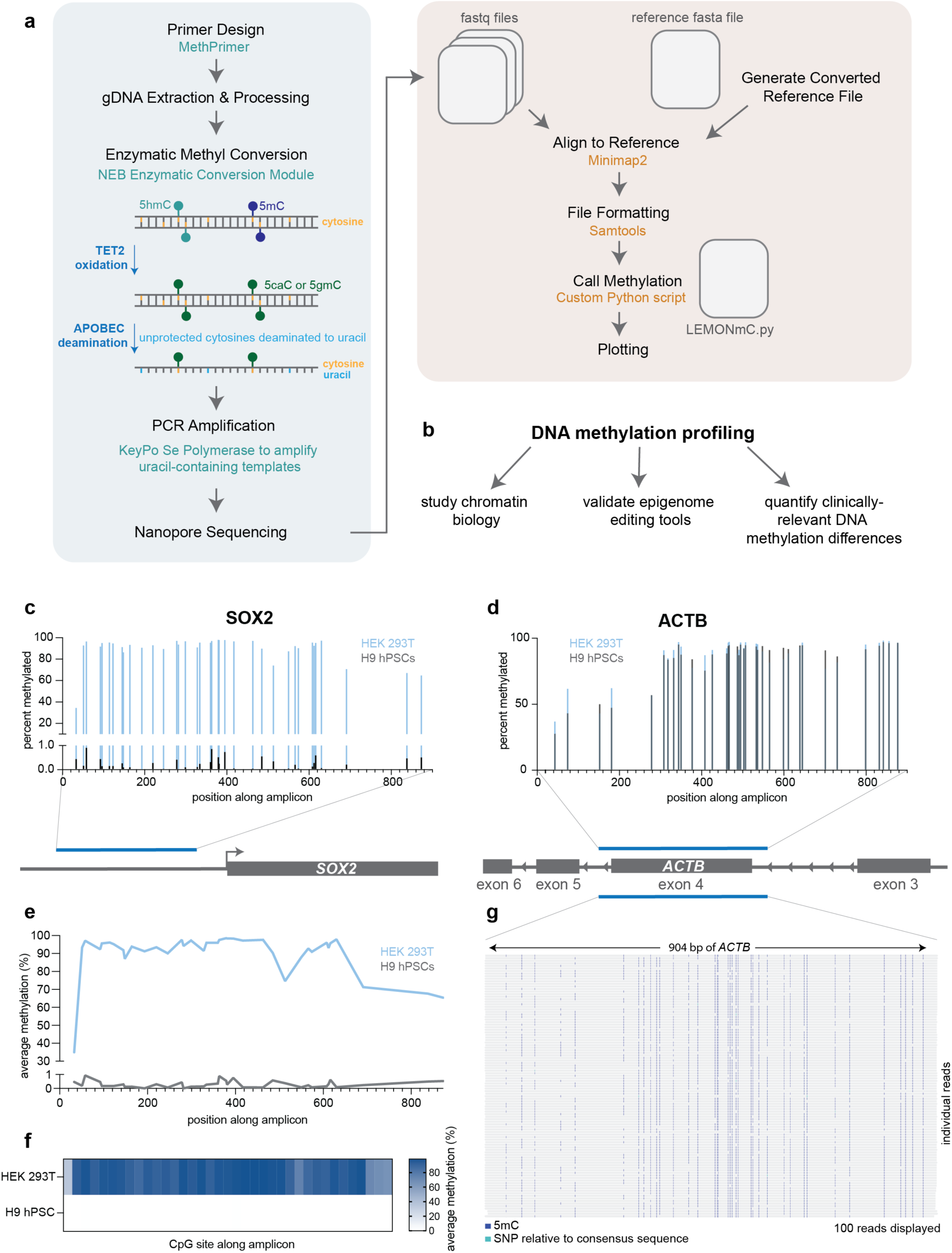
LEMONmethyl-seq for locus-specific, long-read DNA methylation profiling. (a) Experimental and computational workflow of LEMONmethyl-seq. (b) Broad usages of DNA methylation profiling in both fundamental and translational research. (c) Bar plot of LEMONmethyl-seq DNA methylation profiling of the *SOX2* locus in HEK293T (blue) and H9 hPSC (gray) cells. A schematic of the amplicon relative to the *SOX2* gene is shown below. (d) Bar plot of LEMONmethyl-seq DNA methylation profiling of the *ACTB* gene body region in HEK293T (blue) and H9 hPSC (gray) cells. A schematic of amplicon relative to the *ACTB* gene is shown below. (e) Line plot of *SOX2* LEMONmethyl-seq data in HEK293T (blue) and H9 hPSC (gray) cells. (f) Heatmap of *SOX2* LEMONmethyl-seq data in HEK293T and H9 hPSC cells. (g) Integrated Genomics Viewer (IGV) screenshot of 100 individual reads from LEMONmethyl-seq of *ACTB* region in H9 hPSCs. Methylated cytosines are plotted in dark blue. Other SNPs are plotted in teal.

## RESULTS

### LEMONmethyl-seq: long-read, amplification-based, single-nucleotide detection of DNA methylation

We sought to implement a cost-effective, amplification method to profile DNA methylation of long genomic loci in mammalian cells. We chose EM conversion as it is less damaging to genomic DNA compared to bisulfite treatment, thus allowing us to amplify long sequences [22,28,50]. We first tested a panel of different DNA polymerases that are designed to amplify uracil-containing DNA templates. All the polymerases tested successfully amplified EM-converted lambda DNA. However, only the high-fidelity polymerase KeyPo SE consistently produced desired amplicons of endogenous genomic loci from converted human genomic DNA (gDNA) (**Fig. S1a-c**). We additionally found that digestion of gDNA with methylation-insensitive restriction enzymes cutting outside of the locus-of-interest prior to EM conversion can increase PCR yield (**Fig. S1d, Table S1**). For the rest of our experiments, we utilize the KeyPo SE polymerase which results in consistent, robust, and cost-effective amplification across primer sets on EM-converted DNA.

Following PCR amplification, the amplicons are sequenced using long-read technologies, such as Oxford Nanopore, that can be performed in-house or by commercial long-read sequencing services. Drawing from previous work [51,52], we developed a computational pipeline to determine DNA methylation levels using existing packages for alignment and analysis (**Fig. 1a**). This pipeline does not require heavy computational resources and can easily be executed on a laptop. We validated our pipeline on control, unmethylated lambda DNA and observe low false positive rates (**Fig. S1e**). We term this robust, cost-effective targeted DNA methylation profiling method <u>L</u>ocus-<u>E</u>nriched <u>M</u>apping <u>O</u>f <u>N</u>ucleotide methylation sequencing (LEMONmethyl-seq).

We first applied LEMONmethyl-seq to compare DNA methylation at known differentially methylated endogenous genomic sites between two different human cell lines: HEK293T (embryonic kidney) and H9 hPSC (primed pluripotent stem cells). We performed LEMONmethyl-seq of the promoter of *SOX2*, a pluripotency-related gene, by amplifying 868 bases that consists of 37 CpG sites. *SOX2* is expressed highly in stem cells compared to differentiated cells [53]. Indeed, LEMONmethyl-seq of the *SOX2* promoter shows low DNA methylation levels in H9 cells and high in HEK293T cells, with many CpG sites resulting in nearly 100% methylation levels in HEK293T cells (**Fig. 1c**). In comparison, LEMONmethyl-seq of a 904 bp gene body region of *ACTB*, a housekeeping gene, is highly methylated in both lines, consistent with high DNA methylation levels in gene bodies of expressed genes (**Fig. 1d**) [54,55]. These data confirm expected DNA methylation trends and provide site-specific methylation information. The DNA methylation data can be visualized in bar plots highlighting DNA methylation of individual sites and position along the amplicon (**Fig. 1c-d**), line graphs to showcase methylation trends across the amplicon (**Fig. 1e**), heatmaps to contrast differences at methylated sites between samples (**Fig. 1f**), or as single reads to display individual-molecule methylation patterns (**Fig. 1g**).

### Benchmarking LEMONmethyl-seq with bisulfite-PCR and whole-genome EM-seq

To benchmark LEMONmethyl-seq, we started our comparison with bisulfite conversion followed by targeted PCR and single colony picking of amplicons (bisulfite-PCR). We chose our previous bisulfite-PCR data that measured DNA methylation changes at the endogenous promoter of the *HIST2H2BE* after CRISPRoff epigenome editing (**Fig. S2a**) [56]. CRISPRoff is a synthetic fusion of dCas9 to the *de novo* DNA methyltransferase DNMT3A-DNMT3L and the KRAB domain that together establishes DNA methylation and repressive H3K9me3 at targeted gene promoters, resulting in long-term transcriptional repression by epigenetic silencing (**Fig. 2a**) [56].

**Figure 2.**
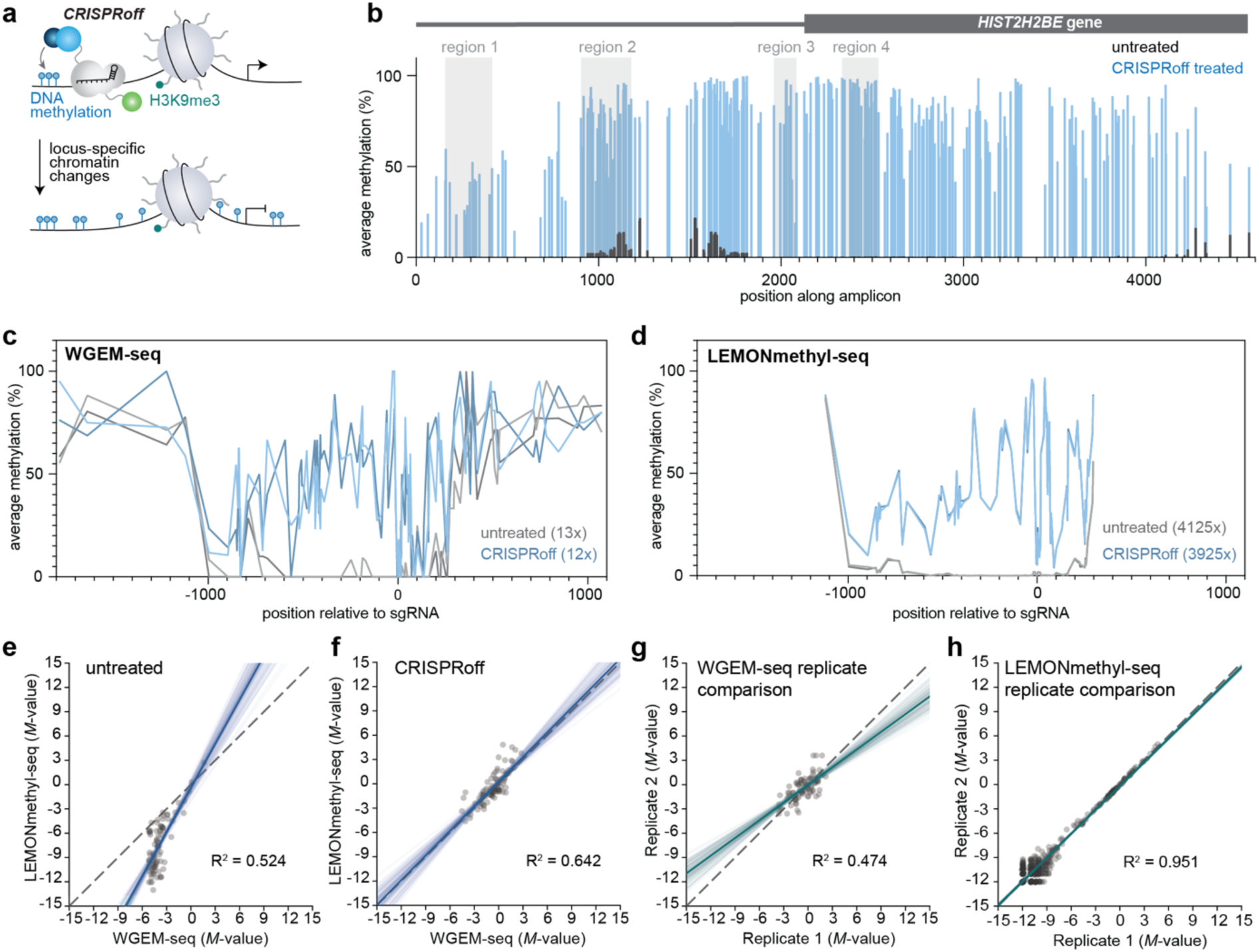
Benchmarking LEMONmethyl-seq to bisulfite-PCR and WGEM-seq. (a) Schematic of CRISPRoff epigenome editing. CRISPRoff leads to the establishment of DNA methylation through the direct fusion of DNMT3A-3L to dCas9. The KRAB domain recruits factors to mediate the establishment of H3K9me3. (b) LEMONmethyl-seq profiling across the *H2B* locus. DNA methylation of CRISPRoff-treated cells is displayed in blue and untreated cells in gray. Bisulfite amplicons (region 1, 2, 3, and 4) from Nuñez et al. (2021) are displayed with light gray boxes. (c) *CD55* DNA methylation profiles from WGEM-seq data from Xu, Besselink, et al. (2025) in Jurkat cells. Two biological replicates of untreated samples are plotted in gray and two CRISPRoff-treated biological replicates are displayed in blue. (d) *CD55* DNA methylation profiles from LEMONmethyl-seq performed on the same gDNA samples as in (c). Two biological replicates of untreated samples are plotted in gray and two CRISPRoff-treated biological replicates are displayed in blue. (e) Correlation of *M*-values of each CpG site from *CD55* amplicon determined by WGEM-seq and LEMONmethyl-seq in untreated Jurkat cells. (f) Correlation of *M*-values of each CpG site from *CD55* amplicon determined by WGEM-seq and LEMONmethyl-seq in CRISPRoff-treated Jurkat cells. (g) Correlation of *M*-values of each CpG site from *CD55* amplicon between each CRISPRoff-treated biological replicate profiled by WGEM-seq. (h) Correlation of *M*-values of each CpG site from *CD55* amplicon between each CRISPRoff-treated biological replicate profiled by LEMONmethyl-seq.

Previously, we used bisulfite-PCR to amplify four distinct regions across the *HIST2H2BE* promoter 30 days after CRISPRoff treatment, each consisting of a ∼200-300 bp amplicon (**Fig. 2b, Fig. S2a-b**) [56]. Using the same cells that generated the bisulfite sequencing data, we performed LEMONmethyl-seq to amplify a single ∼4.5 kb fragment that spans the *HIST2H2BE* promoter and the four regions amplified with bisulfite-PCR (**Fig. 2b**). Comparison of the four regions with LEMONmethyl-seq and bisulfite-PCR amplicons showed similar trends in DNA methylation status across the *HIST2H2BE* locus (**Fig. S2c**). Notably, the bisulfite-PCR datasets consisted of 7-10 Sanger short-sequencing reads, whereas the LEMONmethyl-seq data compiles over 3000 reads from Nanopore sequencing at a lower overall cost.

We next compared LEMONmethyl-seq to whole-genome enzymatic methyl sequencing (WGEM-seq). We chose an existing dataset that used WGEM-seq to profile DNA methylation of the endogenous gene, *CD55*, following CRISPRoff treatment in Jurkat cells (**Fig. 2c**) [57]. Using the same genomic DNA utilized for the WGEM-seq profiling, we performed LEMONmethyl-seq to profile a ∼1.5 kb region around the *CD55* promoter (**Fig. 2d**). LEMONmethyl-seq and WGEM-seq show similar trends of DNA methylation before and after epigenome editor treatment (**Fig. 2e-f**). Notably, LEMONmethyl-seq has over 300x greater sequencing depth across the target region compared to WGEM-seq that was performed at 36x genome coverage.

We next compared DNA methylation detection between methods and biological replicates (**Fig. 2g-h**). We expect no DNA methylation at the *CD55* promoter in untreated samples. Though both methods quantify low DNA methylation of the untreated samples, WGEM-seq has higher methylation averages per site relative to LEMONmethyl-seq, perhaps due to outlier bias with fewer reads (**Fig. 2e**). In the CRISPRoff treated samples, the two methods differ at sites with low DNA methylation levels (**Fig. 2f**), suggesting more continuous classification of DNA methylation with LEMONmethyl-seq. Additionally, LEMONmethyl-seq shows improved consistency between biological replicates compared to WGEM-seq (**Fig. 2g-h, Fig. S2d-e**).

LEMONmethyl-seq recapitulates data obtained from other standard DNA methylation profiling techniques. Furthermore, LEMONmethyl-seq offers improved read length, sequencing depth, and consistency for targeted DNA methylation profiling compared to bisulfite-PCR and WGEM-seq at much lower costs.

### Detection of DNA methylation establishment dynamics following epigenome editing

Epigenome editing has emerged recently as a useful approach for programming gene repression or activation at specified genomic loci. Of note, DNA methylation-based epigenome editors induce long-term heritable transcriptional silencing due to the faithful inheritance of DNA methylation during cell division. Epigenome editing has been applied in cell culture, as well as in mouse and non-human primate models [56–61]. Although the development and clinical use of epigenome editors have matured rapidly, mechanistic understanding of how these technologies remodel the local epigenome of the target site remains limited. Moreover, targeted epigenome editing with DNA methylation often results in propagation of DNA methylation across the targeted locus beyond the sgRNA target site, as exemplified in Fig. 2c with *HIST2H2BE*. However, the mechanism and temporal dynamics of DNA methylation establishment and propagation across the genomic target site remain unclear.

Here, we compared a panel of CRISPR-based epigenome editors with distinct gene silencing dynamics and applied LEMONmethyl-seq to determine the underlying chromatin changes at the target site after epigenome editing. We utilized four CRISPR-based epigenome editors: dCas9 as a control, CRISPRi, CRISPRoff, and CRISPRoff with a point mutation in DNMT3A rendering the enzyme catalytically dead and unable to methylate DNA (CRISPRoff D3A-mut) (**Fig. 3a**). CRISPRi is a fusion of dCas9 to a KRAB domain that recruits chromatin effectors to establish heterochromatic H3K9me3 to induce transcriptional silencing [62,63]. CRISPRoff combines KRAB with the de novo DNMT3A-DNMT3L to program long-term epigenetic silencing by establishing DNA methylation and H3K9me3 that are faithfully inherited through subsequent cell divisions [57].

**Figure 3.**
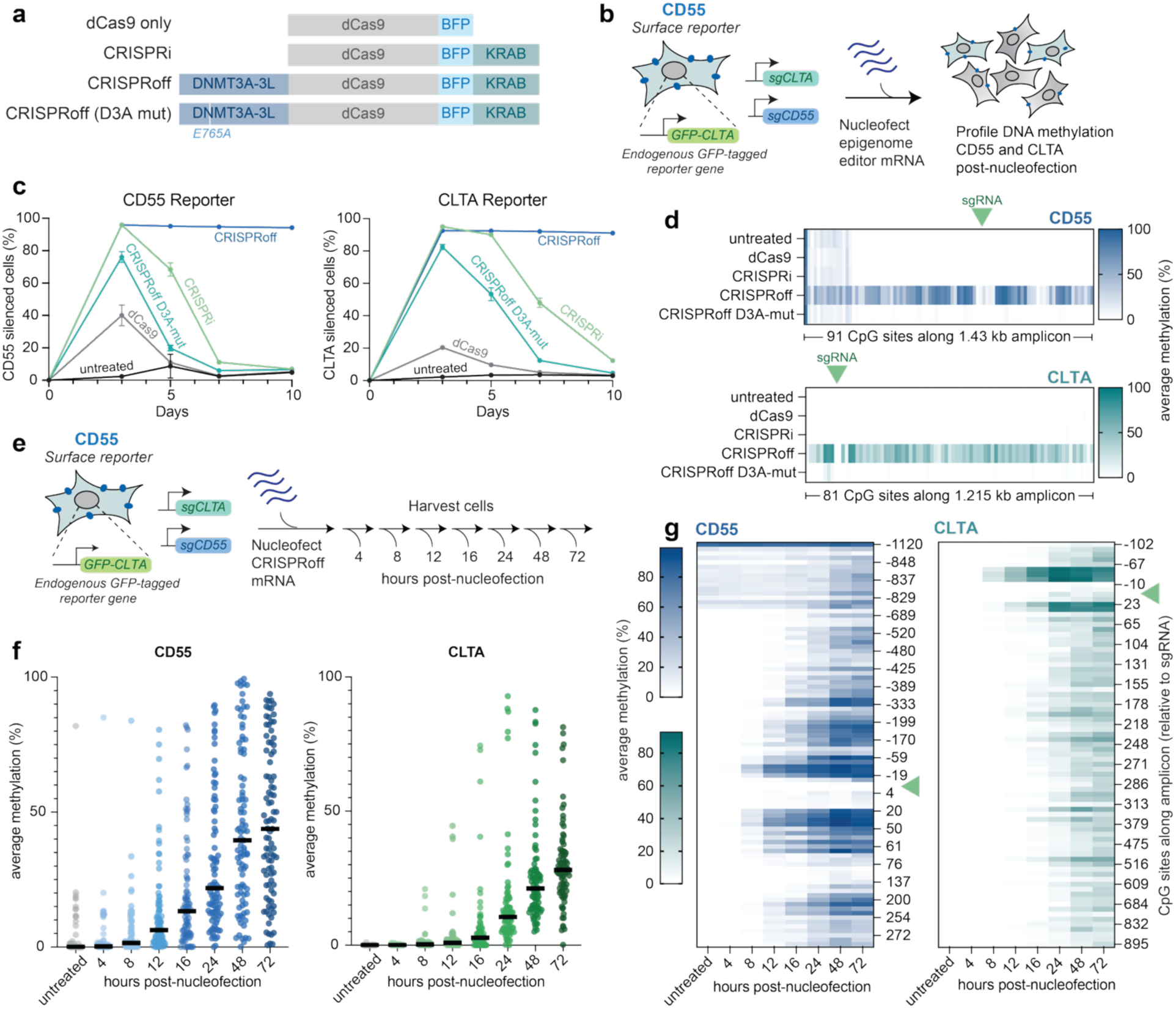
Profiling dynamic DNA methylation establishment following epigenome editing at two endogenous loci. (a) Linear schematics of epigenome editors. (b) Experimental workflow. A HEK293T cell line with *CLTA* endogenously tagged with GFP was engineered to express sgRNAs targeting *CD55* and *CLTA*. Epigenome editor mRNA was introduced to these cells via nucleofection and reporter silencing and DNA methylation establishment were quantified. (c) Quantification of flow cytometry data of *CD55* and *CLTA* silencing at multiple timepoints following epigenome editor nucleofection. Error bars represent standard deviation across three replicates. (d) Heatmaps of LEMONmethyl-seq DNA methylation profiling at day 3 post-nucleofection of different epigenome editors. Only CpG sites are plotted. sgRNA position is annotated with a green arrow. The *CD55* amplicon is 1.43 kb long, spanning 91 CpG sites. The *CLTA* amplicon is 1.215 kb long, covering 81 CpG sites. (e) Experimental workflow. Dual reporter cells were nucleofected with CRISPRoff mRNA and samples were collected at timepoints following nucleofection for DNA methylation profiling. (f) DNA methylation profiling from LEMONmethyl-seq following CRISPRoff mRNA treatment. Each dot represents a single CpG site along the LEMONmethyl-seq amplicon. (g) Heatmaps of average methylation of each CpG site along LEMONmethyl-seq amplicon. Position of CpG site relative to the sgRNA site (green arrow) is annotated.

To assay DNA methylation establishment following epigenome editing, we first engineered a dual guide HEK293T cell line that allows us to concurrently target two endogenous genes in the same cell and compare the dynamics between the two loci for each CRISPR epigenome editor. We utilized a HEK293T cell line with the endogenous *CLTA* gene tagged with GFP to express two distinct sgRNAs, each targeting the promoters of *CD55* and *CLTA*. *CD55* and *CLTA* are nonessential for cell viability, and the CD55 protein localizes to the cell surface that allows us to measure its repression by antibody staining and flow cytometry along with direct readout of CLTA-GFP silencing. To initiate programmable epigenetic repression, we nucleofected mRNAs encoding dCas9, CRISPRi, CRISPRoff, and CRISPRoff D3A-mut into cells and measured CD55 and CLTA silencing at 3, 5, 7, and 10 days post-nucleofection (**Fig. 3b**).

Time course analysis of CD55 and CLTA protein levels show CRISPRoff, CRISPRi, and CRISPRoff D3A-mut induce significant CD55 and CLTA repression at day 3 (**Fig. 3c**). Consistent with previous publications, CD55 and CLTA levels reactivate to the on-state in CRISPRi- and CRISPRoff D3A-mut-treated cells at day 5 and day 7 as they do not have de novo DNA methylation capabilities [56,57,64,65]. In contrast, CRISPRoff-treated cells retain the target genes in the off-state over the 10-day time course due to the faithful maintenance of DNA methylation (**Fig. 3c**).

To couple the protein silencing dynamics with DNA methylation changes, we performed LEMONmethyl-seq in treated cells at 3 days post mRNA delivery. We amplified 1.43 kb across the *CD55* promoter, quantifying DNA methylation of 91 CpG sites, and 1.215 kb across the *CLTA* promoter, capturing DNA methylation of 81 CpG sites. We detect high DNA methylation establishment on the *CLTA* and *CD55* promoters in CRISPRoff-treated cells (**Fig. 3d**). In contrast, no significant DNA methylation is established with dCas9, CRISPRi, or the CRISPRoff D3A-mut at day 3 (**Fig. 3d**). These data highlight that epigenome editors that induce transient silencing of the reporter do not establish DNA methylation, whereas CRISPRoff induces DNA methylation that results in heritable gene silencing. We further profiled the CRISPRoff-treated cells 5 and 10 days post-nucleofection. Consistent with protein-level silencing, we observe maintenance of DNA methylation after CRISPRoff treatment at both *CD55* and *CLTA* (**Fig. S3a**).

Our data show that CRISPRoff-induced DNA methylation is established by day 3 post-nucleofection. Thus, we sought to capture DNA methylation changes at earlier time points to gain insights into the dynamics of establishment and propagation across the target sites. We repeated the CRISPRoff mRNA nucleofection into HEK293T with *CLTA* and *CD55* targets and collected cells for LEMONmethyl-seq at 4, 8, 12, 16, 24, 48, and 72 hours after nucleofection **(Fig. 3e**).

Our LEMONmethyl-seq data reveal gradual DNA methylation establishment during the time course. We observe specific CpG dinucleotides gaining methylation at 8 hours post-nucleofection. By 48 hours, we observe high levels of DNA methylation establishment across the amplicon, which is retained at 72 hours post-nucleofection (**Fig. 3f**). At the protein level, the repression dynamics for CD55 silencing are more rapid than CLTA (**Fig. S3b**), and this pattern is consistent with more gradual DNA methylation establishment at the *CLTA* promoter (**Fig. 3f**).

Comparison of each CpG site across the locus reveals initial establishment around the sgRNA binding site that propagates across the promoter. The average methylation level of these sgRNA-proximal sites increases drastically by 12 hours post-nucleofection and reaches above 90% by 48 hours post-nucleofection at both reporters (**Fig. 3g**). Interestingly, we observe a depletion of DNA methylation directly at the sgRNA site. dCas9 has a footprint of about 78 bp with a strong interaction footprint of 20 bp across the guide RNA site [66–69]. This size correlates with the DNA methylation *glen*, a small genomic region devoid of DNA methylation, where the epigenome editor was originally targeted.

Together, LEMONmethyl-seq reveals rapid DNA methylation establishment and expansion following CRISPRoff treatment and showcases how the method can be applied to study DNA methylation in a dynamic context.

### DNA methylation profiling across transposable element families in static and dynamic cellular contexts

Next, we sought to profile endogenous genomic loci with variable and dynamic DNA methylation patterns. Transposable elements (TEs) are highly abundant in the human genome and are often transcriptionally silenced by DNA methylation [70–76]. We designed primers targeting the consensus regions of the youngest and most active TEs in the human genome: LINE-1 (L1Hs) and HERVK [77–79]. Our L1Hs primer set amplifies a consensus region of the L1Hs 5’ untranslated region (5’ UTR), which functions as an internal promoter to initiate transcription [80,81]. The consensus primers allow us to amplify across many L1Hs genomic loci that will capture global L1Hs DNA methylation trends. Comparison of the individual long reads (100 reads plotted) shows sequence and DNA methylation heterogeneity across the L1Hs 5’ UTR in hTERT-RPE1 cells with notable conserved DNA methylation sites at specific positions (**Fig. 4a**).

**Figure 4.**
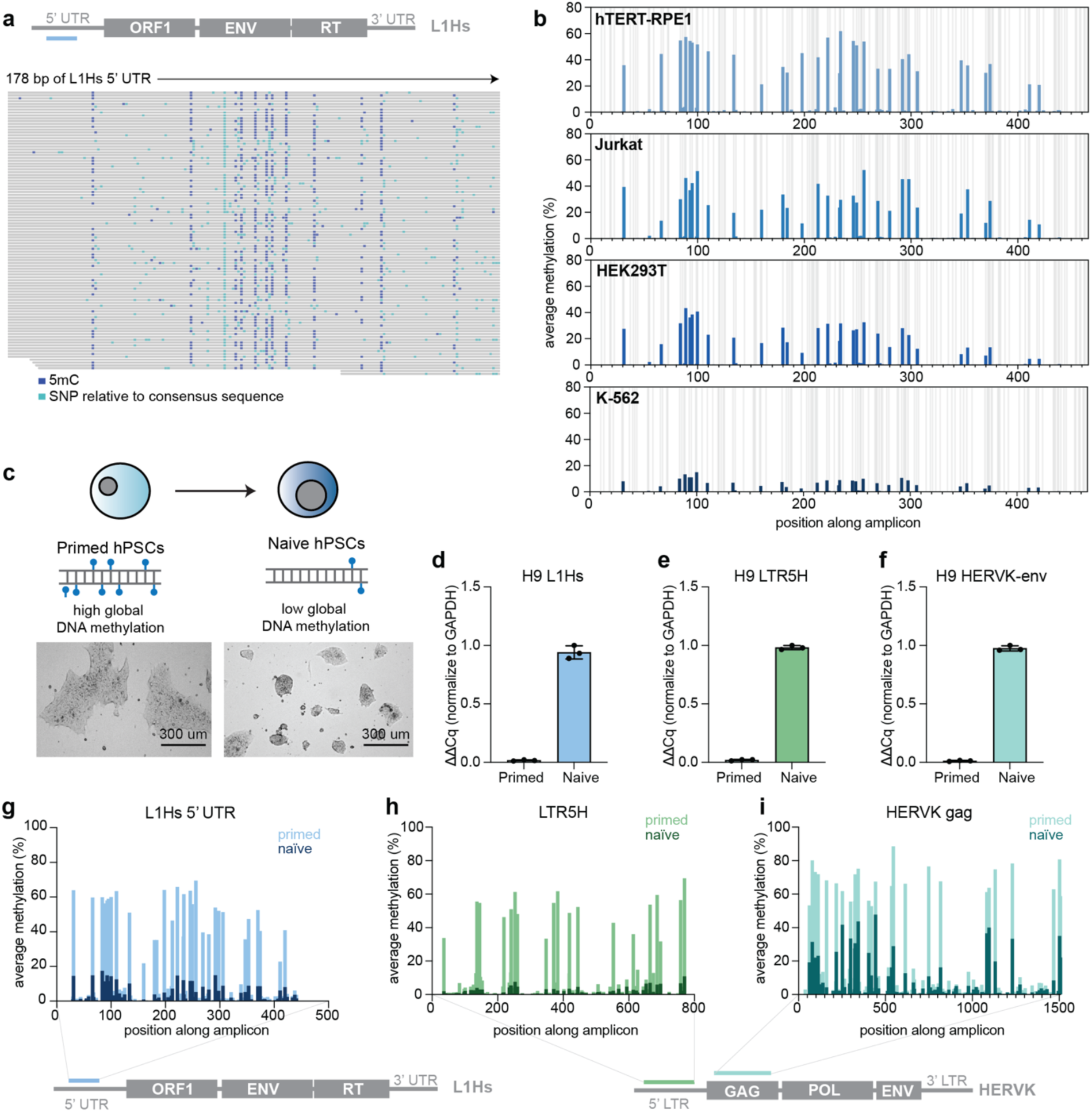
Quantifying DNA methylation trends across transposable element loci. (a) Schematic of L1Hs element and the location of the L1Hs 5’ UTR consensus amplicon. Below, IGV plots of 100 individual reads from hTERT-RPE1 covering the first 178 bp of the L1Hs amplicon. Methylated cytosines are displayed in blue. SNPs relative to the consensus sequence are plotted in teal. (b) LEMONmethyl-seq across L1Hs 5’ UTR in different cell lines (hTERT-RPE1, Jurkat, HEK293T, K-562). Gray bars represent cytosine locations and blue bars represent average methylation at each cytosine site. (c) Schematic of primed to naïve pluripotency induction process along with representative microscopy images of cells in each state. (d-f) RT-qPCR data of (d) L1Hs, (e) LTR5H, and (f) HERVK transcript levels in primed compared to naïve H9 hPSCs. TE transcripts were normalized to GAPDH using the ΔΔCq method and plotted as fold increase relative to the naïve state. Error bars represent standard deviation between three technical replicates. (g-i) LEMONmethyl-seq profiles of primed and naïve H9 hPSCs across (g) L1Hs 5’ UTR, (h) LTR5H, and (i) HERVK consensus sequences. Primed hPSC profiles are shown in the lighter color with naïve profiles in the darker color. Schematics of each amplicon’s position along the TEs are shown below.

We next determined the DNA methylation patterns on TEs across cell types by profiling L1Hs 5’ UTR DNA methylation across common laboratory cell lines: hTERT-RPE1 (retinal pigment epithelial), Jurkat (T lymphoblast), HEK293T (embryonic kidney), and K-562 (lymphoblast) cells. We observe large DNA methylation differences across the L1Hs region across cell lines. Due to sequence variation, we analyzed the methylation of all cytosine positions from the reference sequence, rather than only those annotated as in a CpG context. The maximum methylation observed for hTERT-RPE1, Jurkat, HEK293T, and K-562 cells are 61.82%, 52.27%, 43.38%, and 14.95%, respectively (**Fig. 4b**). Across cell lines, the cytosine positions with the highest relative methylation remained consistent, suggesting potential selection for site-specific methylation of TEs.

We next sought to apply LEMONmethyl-seq in a cellular context with dynamic changes in DNA methylation. During mammalian embryogenesis, early embryonic cells undergo global DNA methylation erasure and establishment during pre-implantation and post-implantation stages, respectively [82–84]. TE genomic loci are among the most differentially DNA methylated loci during these processes, and the loss of DNA methylation can lead to transcriptional activation of TEs [82,85]. We utilized previously established protocols that transition human pluripotent stem cells (hPSC) between primed and naïve states, mimicking the global DNA methylation dynamics in early development [86,87]. We first transitioned H9 hPSCs from a primed state (high global DNA methylation) to a naïve state (low global DNA methylation). We performed reverse transcription quantitative PCR (RT-qPCR) on TE transcripts in primed and naïve hPSCs that show high transcriptional upregulation of L1Hs, LTR5H, and HERVK in naïve hPSCs compared to primed cells (**Fig. 4d-f**).

To correlate transcriptional changes with DNA methylation dynamics, we performed LEMONmethyl-seq on L1Hs, LTR5H, and HERVK genomic loci in primed and naïve hPSCs. In primed hPSCs, we observe high DNA methylation levels across the L1Hs 5’ UTR, LTR5H, and HERVK gag regions. Across the L1Hs 5’ UTR region, we observe high correlation between cell lines, especially between primed hPSCs and hTERT-RPE1 cells which have the highest levels of L1Hs methylation relative to the other cell lines (**Fig. S4b**). Across all regions (L1Hs 5’ UTR, LTR5H, and HERVK gag), we observe a drastic decrease in DNA methylation in the naïve state compared to the primed state (**Fig. 4g-i**). The maximum fold decrease for cytosine methylation across the L1Hs 5’ UTR, LTR5H, and HERVK gag regions were 33.72, 35.33, and 15.85, respectively. These data showcase the global DNA methylation landscape changes during the primed to naïve hPSC transition which drive transcriptional changes and altered cell programming. These data highlight the use of LEMONmethyl-seq to assay TE DNA methylation trends in both static and dynamic systems.

### LEMONmethyl-seq detects endogenous CpH methylation

Canonical DNA methylation in human somatic cells occurs in a CpG context, allowing for methylation inheritance across cell divisions (**Fig. 5a**). In mammalian genomes, CpH methylation (CpA, CpT, CpC) is generally rare, but is enriched in specific biological contexts including neurons, oocytes, and embryonic stem cells [44–49]. Although current DNA methylation profiling methods can capture CpH methylation, the low frequency of this modification requires high sequencing depth to reliably detect and quantify CpH methylation status.

**Figure 5.**
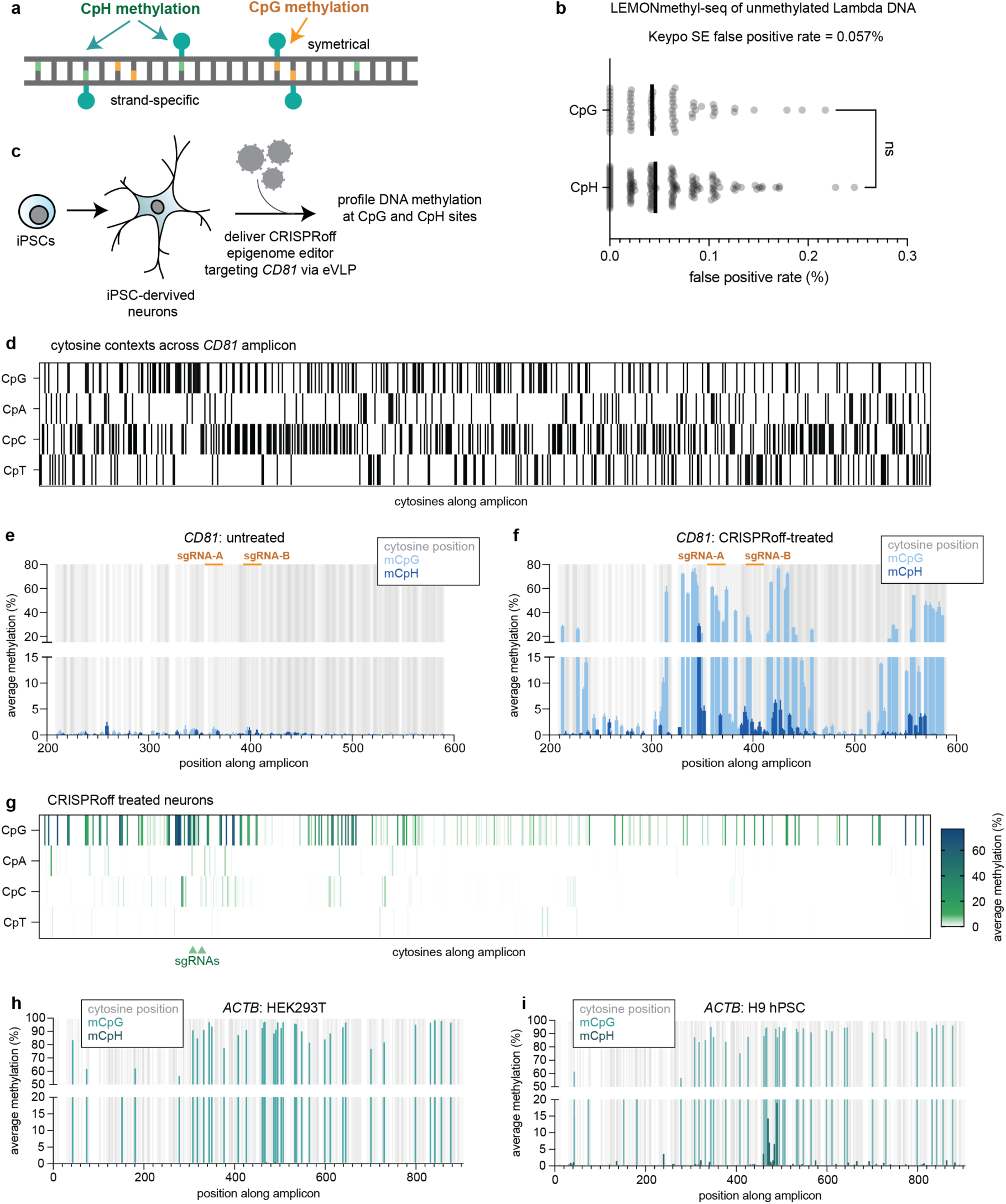
LEMONmethyl-seq for profiling CpH methylation across target loci. (a) Schematic of CpG and CpH methylation. (b) False positive rate of LEMONmethyl-seq between CpG and CpH contexts. LEMONmethyl-seq workflow was performed on control, unmethylated lambda DNA using lambda primer set #3. Any methylation calling corresponds to a false positive. Significance comparison performed using Mann-Whitney unpaired, nonparametric U-test. (c) Experimental workflow of iPSC to neuron differentiation and CRISPRoff epigenome editing at *CD81* through VLP delivery. (d) Location of cytosines in different contexts (CpG, CpA, CpC, and CpT) across the *CD81* amplicon. (e) Bar plot of CpG (light blue) and CpH (dark blue) methylation along *CD81* amplicon in untreated neurons. Cytosine positions are displayed in gray. sgRNA sites are highlighted in orange. Error bars represent standard deviation between two biological replicates. (f) as in (e) but in CRISPRoff-treated neurons. (g) Heatmap of average percent methylation following CRISPRoff treatment of neurons targeted to *CD81*. Cytosine sites are differentiated by context (CpG, CpA, CpC, CpT). sgRNA positions are annotated by green arrows. (h) Bar plot of *ACTB* gene body methylation in HEK293Ts. All cytosine positions are annotated in gray, CpG methylation is plotted in teal, and CpH methylation is plotted in dark teal. (i) as in (h) but in H9 hPSCs.

We first verified low false positive rates for methylation calling in both CpG and CpH contexts. We applied the LEMONmethyl-seq workflow to unmethylated lambda DNA with no DNA methylation in CpG or CpH contexts, thus labeling any detected methylation as a false positive. Across the lambda DNA amplicon, false positive methylation was rare, with a false positive rate of 0.057% across all contexts (**Fig. 5b**). Additionally, the false positive rate did not differ significantly between CpG or CpH contexts (**Fig. 5b**). These data suggest a low false positive rate for LEMONmethyl-seq and its applicability to quantify CpH methylation.

Having established low false positive rates, we next applied LEMONmethyl-seq to assay CpH methylation using CRISPR epigenome editing to detect new CpH methylation at a target locus. We differentiated induced pluripotent stem cells (iPSCs) into neurons and treated the neurons with engineered virus-like particles (eVLPs) to deliver CRISPRoff complexed with a mix of two sgRNAs targeting the promoter of *CD81*, a non-essential gene (**Fig. 5c**). We assayed DNA methylation across a ∼2.1 kb region covering the *CD81* promoter and including 178 CpG sites, 105 CpA sites, 278 CpC sites, and 141 CpT sites (**Fig. 5d**). In the untreated sample, DNA methylation is low, with no CpH sites having over 3% average methylation across reads (**Fig. 5e**).

Following CRISPRoff treatment in neurons, however, we observe site-specific CpH methylation around the guide RNA sites. One CpC site proximal to a sgRNA is methylated in over 25% of the reads. Furthermore, we observe CpH methylation at other regions along the amplicon, especially near sites of CpG methylation. Across the amplicon in CRISPRoff treated neurons, 19 CpH sites are methylated in more than 3% of the reads (**Fig. 5f-g**). We additionally sought to determine the CpH dinucleotide preference across the target site. Averaged across cytosine contexts, CpG sites had the highest average methylation, followed by CpA, CpC, and CpT sites (**Fig. 5g, Fig. S5a-b**). Intriguingly, the same CpH sites are preferentially methylated across biological replicates, perhaps due to proximity to methylated CpG sites or sequence motifs favored by DNA methyltransferases.

To verify that CpH methylation calling was not due to incomplete conversion in highly methylated regions, we analyzed CpH methylation across the *ACTB* amplicon in HEK293T and H9 hPSC cells (**Fig. 1d**). Previous studies have observed CpH methylation in hPSCs, particularly in gene bodies of actively transcribed genes as a result of DNMT3B activity [48]. Given these published data, we expected low levels of CpH methylation in HEK293T cells and higher levels in hPSCs along the *ACTB* amplicon. In HEK293T cells, we do not observe CpH sites with greater than 1% methylation, even though this region is highly methylated at CpG sites (**Fig. 5h**). In contrast, our *ACTB* region in H9 hPSCs had 20 CpH sites with greater than 1% average methylation and two CpH sites with over 14% average methylation (**Fig. 5i**). These data showcase how the increased sequencing depth provided by LEMONmethyl-seq can reveal unique site-specific CpH methylation patterns.

### DNA methylation profiling across *MGMT*, a clinically-relevant gene for glioblastoma

DNA methylation at specific genomic loci is used as a biomarker for various diseases [88]. In glioblastoma, DNA methylation at the promoter of the O-6-methylguanine-DNA methyltransferase (*MGMT*) gene is predictive for patient sensitivity to the chemotherapeutic agent temozolomide (TMZ) as well as prognostic for improved survival [89,90]. Patients with high promoter *MGMT* methylation typically have lower MGMT protein levels and thus are more likely to respond better to TMZ. In clinical settings, short-read bisulfite-PCR is a gold standard Clinical Laboratory Improvement Amendments (CLIA) approved assay for DNA methylation profiling of the *MGMT* promoter [91]. We next applied LEMONmethyl-seq to profile long amplicons of the *MGMT* promoter that allows us to capture the full methylation status and variation across various human glioblastoma cell lines.

We first performed a TMZ sensitivity assay across five glioblastoma cell lines. Based on the cell viability curves, two cell lines (GBM43 and GBM6) are TMZ-sensitive and three cell lines are TMZ-resistant (T98G, SF7996, and LN18) (**Fig. 6a**). To correlate the TMZ response with *MGMT* methylation, we performed LEMONmethyl-seq on each sample by amplifying a 1.6 kb region spanning the *MGMT* promoter and start of the coding region. Across the *MGMT* locus, we detect higher average methylation in the two TMZ-sensitive lines compared to the three resistant cell lines (**Fig. 6b-c**).

**Figure 6.**
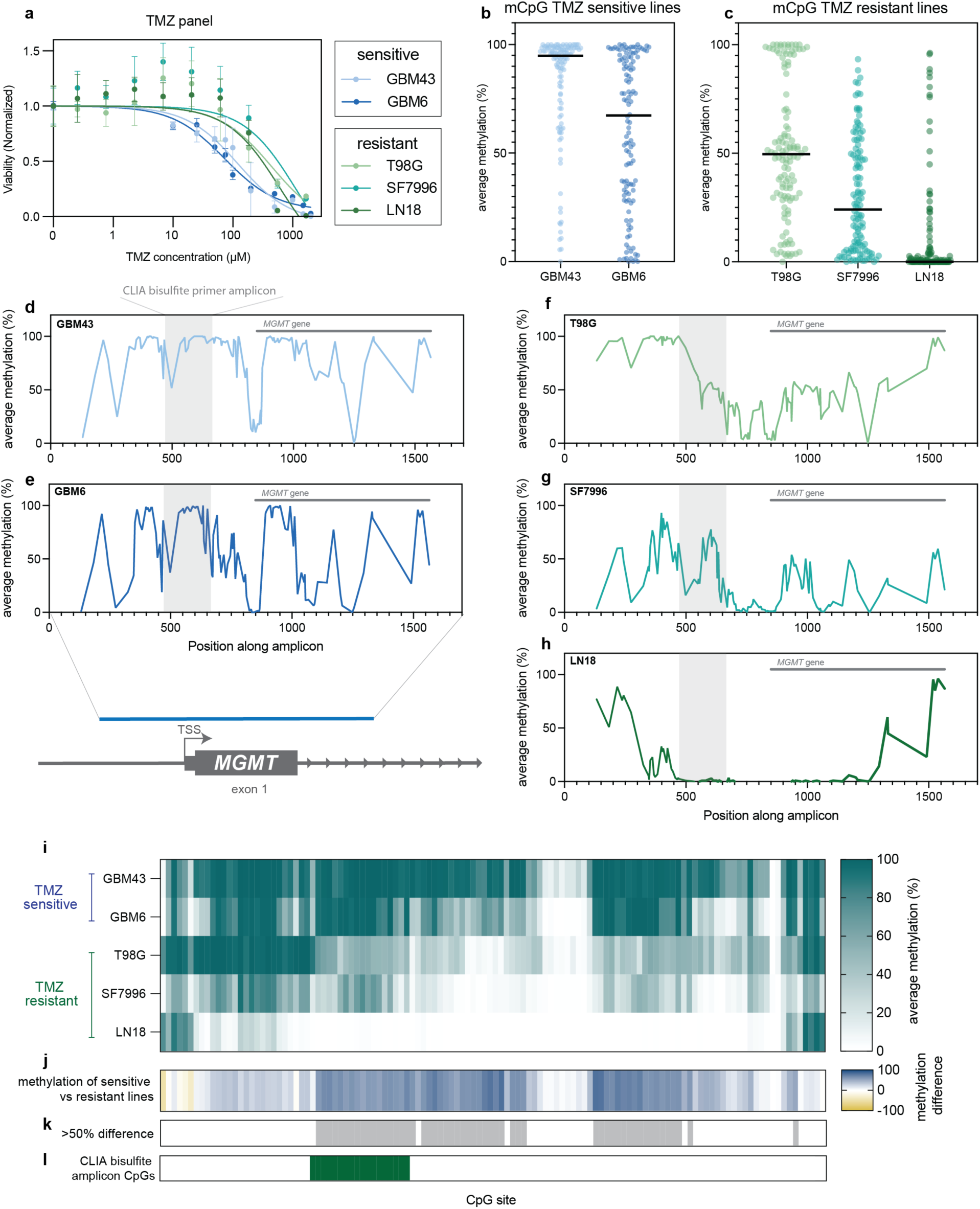
DNA methylation differences across glioblastoma cell lines at the MGMT locus. (a) Cell viability following treatment with TMZ to measure drug sensitivity across glioblastoma cell lines. Data was obtained from 6 technical replicates. The lines plot the results of nonlinear regression. Error bars represent standard deviation across replicates. (b) Average DNA methylation of individual CpG sites across the *MGMT* amplicon in TMZ-sensitive glioblastoma cell lines. (c) As in (b) but for TMZ-resistant glioblastoma lines. (d-h) Average CpG methylation across the *MGMT* amplicon for each glioblastoma cell line. Shaded gray region represents amplicon generated using clinical bisulfite primers. (i) Heatmap of *MGMT* DNA methylation levels across glioblastoma cell lines. (j) Difference between average methylation of TMZ-sensitive and TMZ-resistant lines at each CpG site. Positive values indicate higher average methylation in TMZ-sensitive lines (plotted in blue); negative values indicate higher average methylation in TMZ-resistant lines (plotted in yellow). (k) CpG sites with greater than 50% difference in average DNA methylation between sensitive and resistant lines methylation are plotted in gray, highlighting regions with the largest difference between phenotypic groups. (l) CpG sites covered by the CLIA bisulfite primer set are highlighted in green.

DNA methylation patterns across the locus differ between glioblastoma cell lines and have unique patterns beyond the clinically approved bisulfite-PCR amplicon (**Fig. 6d-i**) [91]. We quantified the difference in average methylation of the TMZ-sensitive cell lines (GBM43 and GBM6) and the TMZ-resistant cell lines (T98G, SF7996, and LN18) along the *MGMT* locus. As expected, the majority of CpG sites showed higher methylation in the sensitive lines compared to the resistant lines (**Fig. 6j**). We further annotated sites with a difference greater than 50% difference between the TMZ-sensitive and resistant lines (**Fig. 6k**). In addition to differential methylation at the CLIA-assayed location, LEMONmethyl-seq revealed high differential methylation immediately downstream the TSS between TMZ-sensitive and TMZ-resistant GBM cell lines. This comparison quantifies methylation across 120 CpG sites spanning a region >8 times larger than the CLIA-assayed locus, which may provide useful, clinically-relevant information about drug sensitivity and epigenome editing that would not have been captured on a gold standard clinical assay [92].

## DISCUSSION

Here we present LEMONmethyl-seq, a method for long-read, targeted DNA methylation profiling paired with a streamlined, computationally-light analysis pipeline. We demonstrate this method to be robust across experimental applications, including DNA methylation profiling following epigenome editing, across TE loci, and at a clinically-relevant gene.

Previous groups have performed EM conversion followed by PCR amplification and second- or third-generation sequencing [28–30], though this workflow has not been widely adopted due to PCR amplification challenges and requirements for in-house sequencing and computational equipment. To develop LEMONmethyl-seq, we first tested commercially available DNA polymerases to efficiently amplify uracil-containing templates, ultimately choosing the KeyPo SE polymerase that consistently yields the desired PCR amplicons. We further streamlined the analysis pipeline such that the computational analysis can easily be performed on a laptop.

Current DNA methylation profiling methods face additional limitations due to false positive methylation calling. False positives in bisulfite-PCR result from incomplete conversion [93], and Nanopore sequencing has a high error rate of methylation calling, especially in CpH contexts [41–43]. We find the LEMONmethyl-seq pipeline with KeyPo SE has a low false positive rate across cytosine contexts. Additionally, with LEMONmethyl-seq, we achieve high read depth (∼3000x) across target loci, orders of magnitude above other DNA methylation profiling methods such as bisulfite-PCR and WGEM-seq. LEMONmethyl-seq’s low false positive rate, in addition to high read depth, underscores its ability to capture subtle DNA methylation dynamics and CpH methylation.

LEMONmethyl-seq is cost-effective compared to other locus-specific DNA methylation profiling methods. Bisulfite-PCR is the most commonly used method for targeted DNA methylation profiling. Our calculation of bisulfite-PCR with 10 Sanger reads costs 193% more than performing LEMONmethyl-seq with over 3000 reads, not including undesired Sanger reads that typically arise from bisulfite-PCR experiments (**Table S3**). Additionally, LEMONmethyl-seq has the ability to profile DNA methylation of much longer regions, providing additional data at a fraction of the cost.

Our work showcases the broad applicability of LEMONmethyl-seq across different experiments. We observe DNA methylation establishment dynamics following epigenome editing. We detect initial DNA establishment directly adjacent to the sgRNA binding site followed by expansive establishment at later timepoints. Interestingly, the DNA methylation establishment profiles following CRISPRoff treatment are similar across cell types and delivery modalities, suggesting relative distance from the epigenome editor and DNA sequence motifs may contribute to establishment patterns (**Fig. S3c**).

We also highlight LEMONmethyl-seq in profiling CpH methylation with high precision. Across LEMONmethyl-seq amplicons in both neurons and hPSCs, we observe relatively high levels of site-specific CpH methylation compared to other cell types. Across the *CD81* amplicon following epigenome editing, the CpH motif with the highest average methylation is CAC, consistent with previously observed CpH motif preferences in neurons [48]. In hPSCs, previous work has identified CAG sites as the preferred CpH methylation motif, especially in the gene bodies of actively transcribed genes due to DNMT3B activity [48,94,95]. Across our *ACTB* amplicon, we find the most highly methylated CpH motifs are CCG, CAG, and CTA. Further studies across additional loci can probe CpH motif preferences in addition to the biological significance of this epigenetic modification.

We note several current limitations and future improvements of LEMONmethyl-seq. Inherently, LEMONmethyl-seq profiling is limited by the size of the amplicon. Additionally, the methylation calling pipeline relies on an accurate reference sequence to determine cytosine positions and cytosine context (CpG versus CpH). In cases of sequence variants (such as across TE loci), we recommend analyzing methylation at all cytosine positions, regardless of context. Additionally, LEMONmethyl-seq as currently described only provides single-stranded DNA methylation information. Strand-specific DNA methylation analysis may be important for certain experiments, especially when profiling the nonsymmetrical CpH methylation landscape. Similar to bisulfite methods, two primer sets, one for each strand, can be designed to quantify strand-specific DNA methylation.

## CONCLUSIONS

In summary, we present <u>L</u>ocus-<u>E</u>nriched <u>M</u>apping <u>O</u>f <u>N</u>ucleotide methylation (LEMONmethyl-seq): an optimized, cost-effective pipeline for single-nucleotide, targeted DNA methylation profiling. Our results reveal patterns and dynamics of DNA methylation following epigenome editing, across transposable elements, and at a clinically-relevant gene. Broadly, our results showcase the wide applicability and usage of LEMONmethyl-seq across experimental contexts. Altogether, we envision future applications of LEMONmethyl-seq in studying fundamental chromatin biology, diagnostics, and translational research.

## METHODS

### LEMON-seq primer design

Primers were designed to amplify EM converted genomic DNA using MethPrimer (https://www.methprimer.com/) [96]. Default parameters were used, except ‘product size’ was adjusted to generate longer amplicons. See Table S1 for oligos used for each locus. For primers amplifying single genomic loci, reference sequences from hg38 (GRCh38 EnsemblRelease 104) were used [97]. For TE primer design, TE consensus sequences were obtained from RepBase for L1HS (L1, Homo sapiens), LTR5_Hs (ERV2, Primates), HERVK (ERV2, Primates). See Table S5 for complete consensus sequences [98]. Dfam was used to further annotate the sequences amplified by our TE consensus primers [99].

### gDNA extraction and optional DNA digestion

Genomic DNA was extracted from 500,000 to 5 million cells using isopropanol precipitation. In short, cells were pelleted and washed with 1x PBS. Cells were resuspended in 500 μL lysis buffer (100 mM Tris-HCl pH 8, 5 mM EDTA, 0.2% SDS, 200 mM NaCl, 100 μg Proteinase K/mL, 100 μg RNase A/mL). Cells were incubated at 55°C with continuous agitation for 1 hour. Samples were centrifuged at room temperature for 15 minutes at max speed and supernatant was added to ice cold isopropanol. The mixture was vortexed and spun at 4°C for 10 minutes at max speed. Isopropanol was removed and the DNA pellet was washed with 70% ice cold ethanol. After another spin at 4°C for 10 minutes at max speed, ethanol was removed and pellet was left to dry for 3 minutes or until ethanol was not visible. DNA was resuspended in 50 μL - 300 μL of nuclease-free water.

When applicable, gDNA was digested with restriction enzymes (NEB). See Table S1 for restriction enzymes and buffers used. For each digestion, 1 μg of gDNA was mixed with buffer and restriction enzyme(s). The mixture was incubated at optimal enzyme temperature for 1 hour. Digested gDNA was purified with DNA purification columns (Qiagen, 28104) and eluted in nuclease-free water.

### Enzymatic methyl conversion

200 ng of gDNA (digested or undigested) was converted with the NEBNext® Enzymatic Methyl-seq Conversion Module (NEB, E7125S/L) using formamide denaturation according to manufacturer’s instructions.

### PCR amplification of EM-converted DNA

PCR amplification of EM-converted DNA was performed with 2 × KeyPo SE Master Mix (Vazyme, PK510-01) according to the manufacturer’s instructions. In short, a 50 μL total PCR reaction was prepared: 20 μL of EM-converted DNA (one EM reaction), 1.5 μL of each 10 μM primer, 25 μL KeyPo SE, and 2 μL water. For the PCR, a 60°C annealing temperature, an extension time of one minute per kb, and 35 cycles were used. After amplification, 5 μL of the PCR reaction was run on a gel to verify successful amplification. Amplified DNA was then purified with DNA purification columns (Qiagen, 28104) and eluted in nuclease-free water.

### Long-read sequencing

Sequencing was performed using the following long-read sequencing services (UC Berkeley DNA Sequencing Facility or Plasmidsaurus). Oxford Nanopore Sequencing was performed by the UC Berkeley DNA Sequencing Facility with the Rapid Barcoding protocol. Premium PCR Sequencing was performed by Plasmidsaurus using Oxford Nanopore Technology with custom analysis and annotation.

### LEMONmethyl-seq computational analysis

Each LEMONmethyl-seq experiment was accompanied by a computational analysis that produced a table of per-cytosine methylation read percentages and counts as follows. First, a strand-specific “in silico EM-converted” reference genome was made using the sed command to replace C with T in the original FASTA reference sequence. Reads were aligned to this in silico EM-converted reference genome using minimap2 (with -ax map-ont --sam-hit-only) [100]. Due to minimap2’s error tolerance, we did not need to convert cytosines in reads for efficient mapping. Aligned SAM files were converted to BAM files using samtools [101,102], which were also sorted and indexed for ease of downstream viewing and analysis. A custom Python script (using the pysam library [103,104]) was run to determine the methylation read percentages and counts at each CpG and CpH site. For each aligned read, cytosines retained as “C” were classified as methylated, and cytosines converted to “T” were classified as unmethylated. To aid ourselves and others in applying these computational steps, the pipeline is available as building blocks in the Honeybee programming system and Honeybee can be used to generate full LEMONmethyl-seq analysis pipelines (*e.g.,* for Figure 2) [105].

### Plotting and correlation analysis

Average methylation data was plotted in Prism 10 for macOS (GraphPad). Individual reads were visualized in Integrative Genomics Viewer (IGV) version 2.16.0 with the following settings adjusted from the default: squished and downsample reads (100). To change colors to teal for SNPs in TE amplicons, exported IGV SVG files were colored with Adobe Illustrator 2024.

Correlation coefficients between recorded methylation profiles (e.g. between LEMONmethyl-seq results and Whole Genome EM-seq results or between replicates of a single method) were calculated via Pearson’s R^2 on the per-CpG methylation counts converted to *M*-values as described by Du et al. (with α = 1) [106]. Linear regression for the correlation plots was bootstrapped by resampling the data (with replacement) 100 times. Correlation calculation and plotting was performed using polars [107], matplotlib [108], and SciPy [109] in a custom Python script.

### Cell culture

#### HEK293T, RPE, Jurkat, K562

All cell lines were obtained from the UC Berkeley Cell Culture Facility. HEK293T were cultured in DMEM (Gibco 11995073), RPE were cultured in DMEM/F12 (Gibco,11330032), Jurkat and K562 were cultured in RPMI (Gibco, 22400105). All were supplemented with 10% (v/v) FBS (VWR), 100 units/mL streptomycin, 100 μg/mL penicillin, and 2 mM glutamine (Gibco, 10378016). All cell lines were cultured at 37°C with 5% CO_2_ in tissue culture incubators.

#### Primed hPSCs culture

Human Pluripotent Stem Cell line H9 (WA09 Lot #WB68075, WiCell) was cultured in primed pluripotency conditions using mTeSR Plus media (Stem Cell Technologies, 100-0276) and hESC-qualified Matrigel (Corning, 354277)-coated plates following manufacturer’s instructions. Briefly, hESCs-qualified Matrigel was initially pre-diluted in ice-cold DMEM/F12 (Gibco, 10565018) supplemented with 15 mM HEPES (Gibco, 15630080), and plates were incubated at room temperature for an hour before hPSCs were seeded. hPSCs were split every 4-5 days when they were at 70-80% confluence. hPSCs were harvested into cell clumps by incubating them for 3 minutes at 37°C with Gentle Cell Dissociation Reagent (Stem Cell Technologies, 100-0485). Media was refreshed every other day, and cells were cultured at 37°C under normoxia conditions (20% O_2_ and 5% CO_2_).

#### Glioblastoma cell culture

LN18, T98G, and SF7996 glioblastoma cells were cultured in DMEM or DMEM-F12 supplemented with 10% fetal bovine serum (Thermo Scientific, 11965084) and antibiotic-antimycotic (Thermo Scientific, 15240062). GBM6 and GBM43 cells were cultured in Neurobasal-A Medium (Gibco, 10888022) supplemented with B-27 (Gibco, 12587010), N-2 (Gibco, 17502-048), plasmocin (Invivogen, ant-mpp), primocin (Invivogen, ant-pm-1), EGF at 20 ng/mL (VWR, 10781-694), and FGF at 20 ng/mL (PeproTech, 10777-988). All cells were routinely confirmed free of mycoplasma using the MycoAlert PLUS Mycoplasma Detection Kit (Lonza, 75860-362).

### sgRNA design

We utilized the lentiviral pLG1 library vector (Addgene #217306) that encodes for expression of a sgRNA from a modified U6 promoter. Additionally, this plasmid encodes for expression of a puromycin resistance gene and mCherry separated by a T2A cleavage sequence from an EF1a promoter. This plasmid was used for expressing the CLTA-targeting sgRNA.

We used an additional lentiviral vector expressing only HALO, rather than the puromycin resistance gene and mCherry. To generate this plasmid, the gene encoding HALO expression was PCR amplified with primers containing overhangs using KAPA HiFi HotStart ReadyMix PCR Kit (Roche). The original pLG1 plasmid was digested with NheI and EcoRI, and the plasmid was constructed using Gibson assembly with ClonExpress Ultra One Step Cloning Kit V3 (Vazyme, C117-01). This vector was used for expressing the *CD55*-targeting sgRNA.

To replace the protospacer sequences, two complimentary oligos were ordered (IDT) with compatible BstXI and BlpI overhangs and ligated into the lentiviral backbone digested with BstXI and BlpI. sgRNA sequences and oligos can be found in Table S2. Protospacer sequences were chosen based on previous algorithms to predict optimal CRISPRi sgRNAs targeting gene promoters [110].

### Cell line generation

To generate the dual guide HEK293T line, CLTA-GFP HEK293T cells expressing CLTA guide with mCherry selection marker were transduced with lentivirus expressing CD55 targeting guide with the HaloTag selection marker. To produce lentivirus, 500,000 HEK293T cells were plated in a 6-well plate. After 12 hours, the plated cells were transfected with the *CD55* HaloTag guide plasmid alongside standard packaging plasmids and TransIT-LT1 Transfection Reagent (Mirus, MIR2306) according to manufacturer’s instructions. Media was changed about 24 hours post-transfection, and the viral supernatant was harvested after 50 hours. The supernatant was filtered through 0.45 mm PVDF syringe filter and 300 μL was plated with CLTA-GFP HEK293T cells with CLTA guide in a 6-well plate. Post-lentivirus addition, CLTA-GFP HEK293T cells were stained for 30 minutes using Janelia Fluor 646 HaloTag Ligand, washed 3x with PBS, and sorted on BDFACS Aria Fusion sorter for HaloTag positive cells. These cells were then expanded for downstream experiments.

### Human Naïve hPSCs resetting

Primed hPSCs were cultured at least two passages in primed culture conditions as described above. hPSCs were reset to naïve pluripotency following the protocol described in Khan et al,. 2021 [86]. One day before starting the conversion, mitomycin C-inactivated mouse embryonic fibroblast (iMEFs) were seeded in a 6-well plate coated in 0.1% Gelatin (Stem Cell Technologies, 07903) at a density of 10^5^ iMEFs/cm^2^. On the following day, primed hPSCs were harvested at 70% confluency and dissociated into single-cell suspension by using Accutase (Gibco, A1110501). 2x10^5^ hPSCs were seeded in each well with mTeSR Plus supplemented with 10 μM Y-27632 inhibitor (MedChemExpress, HY-10071) at normoxia conditions (20% O_2_ and 5% CO_2_). 48 hours later, the media was switched to PXGGA media (composition described in **Table S4**), and cells were cultured under hypoxia (5% O_2_ and 5% CO_2_) from that day onwards. PXGGA media was changed every other day. Cells were split for the first time after 12 days by incubating them at 37°C with Accutase for 3 minutes quenching the reaction by adding equal amounts of PXGGA media. Cells were then centrifuged at 300 g for 5 minutes at room temperature before seeding and were replated at a 1:6 ratio relative to the original plate. Since the first passage (day 12) onwards, cells were split every 4 days, changing the PXGGA media every other day and supplementing it with 10 μM Y-27632 inhibitor during the first 24 hours. Experiments were performed in naïve hPSCs cultured for at least 5 passages in PXGGA media.

### Differentiation and epigenome-editor treatment of iPSC-derived neurons

Neurons were derived from WTC-NGN2 induced pluripotent stem cells (iPSCs) via a doxycycline-inducible neurogenin-2 (NGN2) protocol, as previously described by Ramadoss et al. 2025 [111]. Briefly, differentiation was initiated on Day 0. On Day 3 of differentiation (Day −14 relative to treatment), cells were seeded at a density of 100,000 cells per well onto 24-well plates coated with poly-D-lysine (PDL). Full medium exchanges were performed on Day 7. To prevent neuronal detachment, half medium exchanges were adopted on Day 10. On Day 17, the culture medium was completely removed and replaced with fresh medium containing RENDER-CRISPRoff particles (added as 20 μL of 100-fold concentrated stock per well). Neurons were harvested 7 days post-RENDER treatment (Day 24). Cells were washed once with phosphate-buffered saline (PBS), pelleted by centrifugation, and stored at −80°C for downstream LEMONmethyl-seq analysis.

### Epigenome editor mRNA preparation

For comparing the epigenome editors (Fig. 3a-d), dCas9, CRISPRi, CRISPRoff, and CRISPRoff D3A-mut mRNA was synthesized in-house using the protocol described previously in Pattali et al., 2025 [64]. Briefly, for the in-house synthesis, the epigenome editor sequence was cloned into the pALD-CV42 [T7] (Aldevron) vector. The resulting plasmid was then digested with XhoI and used as a T7-driven template for in vitro transcription using the mMessage mMACHINE T7 ULTRA kit (Invitrogen, AM1345). Following DNase treatment, transcripts were polyadenylated, purified by lithium chloride precipitation, and resuspended in nuclease-free water. mRNA concentration was measured by Qubit RNA HS assay (Q32852) or a spectrophotometer. Purified mRNA was stored at −80 °C until use.

For the CRISPRoff early timepoint experiment (Fig. 3e-g), CRISPRoff mRNA was purchased from Aldevron.

### Epigenome editor mRNA nucleofection

For the epigenome editor comparison, 200,000 cells were nucleofected with 2 μg of epigenome editor (dCas9, CRISPRi, CRISPRoff, CRISPRoff-D3A-mut) mRNA made in-house. Nucleofection was performed using the SF Cell Line 4D-Nucleofector® X Kit S (Lonza, V4XC-2032) following the manufacturer’s instructions with program CM-130 for HEK293Ts. For the CRISPRoff early timepoint experiment, 1 million cells were nucleofected with 10 μg of CRISPRoff mRNA from Aldevron. Nucleofection was performed with the SF Cell Line 96-well Nucleofector® Kit (Lonza, V4SC-2096) following the manufacturer’s instructions with program CM-130 for HEK293Ts.

### Flow cytometry

Flow cytometry and cell staining was performed as described by Christenson, Divekar, et al. 2025 [65]. In short, protein expression of CD55 was assessed by cell surface antibody staining of live cells. Cells were mixed with diluted APC Human Anti-CD55 antibody (BioLengend, 311312) and incubated in the dark at room temperature for 30 minutes. Cells were then washed once with PBS and protein levels were measured on the BD FACSymphony A1 Cell Analyzer (BD Biosciences). For GFP-tagged HEK293T reporter lines, GFP expression was directly measured on the flow cytometer. Data analysis was performed using FlowJo (v10.10).

### Reverse transcription quantitative PCR

Two 6-well plates of naïve and primed hPSC cells were lysed in three volumes of NP-40 lysis buffer (50 mM HEPES-KOH pH 7.5, 150 mM KCl, 5 mM MgOAc, 0.5% Nonidet P-40 alternative, 0.5 mM DTT). Total RNA was extracted from the lysate using phenol-chloroform extraction with acid phenol:chloroform:isoamyl alcohol (25:24:1, pH 5.2, Fisher Scientific) and ethanol precipitation. The total RNA samples were precipitated overnight at −20°C, then centrifuged for 30 minutes at 21000 rcf at 4°C. RNA pellets were washed with 70% ethanol, air-dried for 2 minutes, and resuspended in nuclease-free water. One microgram of total RNA was treated with RQ1 DNase kit (Promega) for 30 minutes at 37°C and incubated with random hexamers for 5 minutes at 65°C. cDNA was generated from RNA using random hexamers by reverse transcription using SuperScript^TM^ II Reverse Transcriptase kit (Thermo) according to the manufacturer’s instructions. Quantitative PCR was performed using iTaq^TM^ Universal SYBR^R^ Green Supermix (Bio-Rad) on the CFX Opus 96 Real-time PCR system. Gene expression levels of L1HS, LTR5H, and HERVK were normalized to GAPDH using the 2-ΔΔCq method. Primer sequences can be found in Table S2.

### Temozolomide cell viability assays

Glioblastoma cells were seeded at 1000–5000 cells per well in 6 replicates per drug dose on a 96-well plate. TMZ was prepared at 100 mM stock (Sigma-Aldrich, T2577). TMZ was added to cell cultures across 1:300 serial dilutions. Cell Titer Glo cell viability assays were performed per manufacturer’s recommendations (Promega, G7571) 7 days following one time addition of TMZ. Luminescence was measured using GloMax 96 Microplate Luminometer (Promega, E6521). Data values were normalized to the vehicle-only condition.

## DECLARATIONS

### Ethics approval and consent to participate

Not applicable

### Consent for publication

Not applicable

### Availability of data and materials

All DNA sequencing results will be available in the Sequence Read Archive (SRA) of the National Center for Biotechnology Information (NCBI). The LEMONmethyl-seq computational pipeline and corresponding code are available on GitHub (https://github.com/Nunez-Lab) and as reconfigurable building blocks in the Honeybee programming system (https://honeybee-lang.org/) [105].

### Author contributions

A.E.C. and J.K.N. initiated and led the study, designed the experiments, and wrote the manuscript with assistance from the co-authors. A.E.C. performed the initial optimizations and data analysis along with the majority of the LEMONmethyl-seq experiments. A.E.C. and J.P.L. wrote the LEMONmethyl-seq computational analysis pipeline. L.G.P. performed the *ACTB* and *SOX2* profiling. J.P.L. and P.J.C. performed the WGEM benchmarking comparison, and J.P.L. performed the statistical analyses for this comparison under supervision of S.E.C. N.S.D., A.E.C., and P.J.C. performed the epigenome editor comparison experiments. A.E.C with assistance from R.K.P. performed the early timepoint LEMONmethyl-seq profiling following CRISPRoff treatment. R.K.P. and P.J.C. performed the L1Hs 5’ UTR profiling across cell lines experiment. L.G.P. established the primed to naïve induced pluripotency protocol for the lab, N.P. performed RT-qPCR to assay TE expression, N.S.D. performed the LEMONmethyl-seq profiling across L1Hs, LTR5H, and HERVK. B.M. performed the cell culture experiment to differentiate iPSCs to neurons. D.X. performed the LEMONmethyl-seq profiling of the neurons. K.L. and A.H. performed the TMZ sensitivity assay under supervision of S.J.L. A.H. cultured glioblastoma cell lines for LEMONmethyl-seq profiling of the *MGMT* locus.

## Acknowledgements

We thank Joseph McKenna for technical advice and suggestions and Daniella Starr for assisting with the initial transposable element profiling experiments. Additionally, we thank all members of the Nuñez lab for helpful discussions.

---

**Figure S1.**
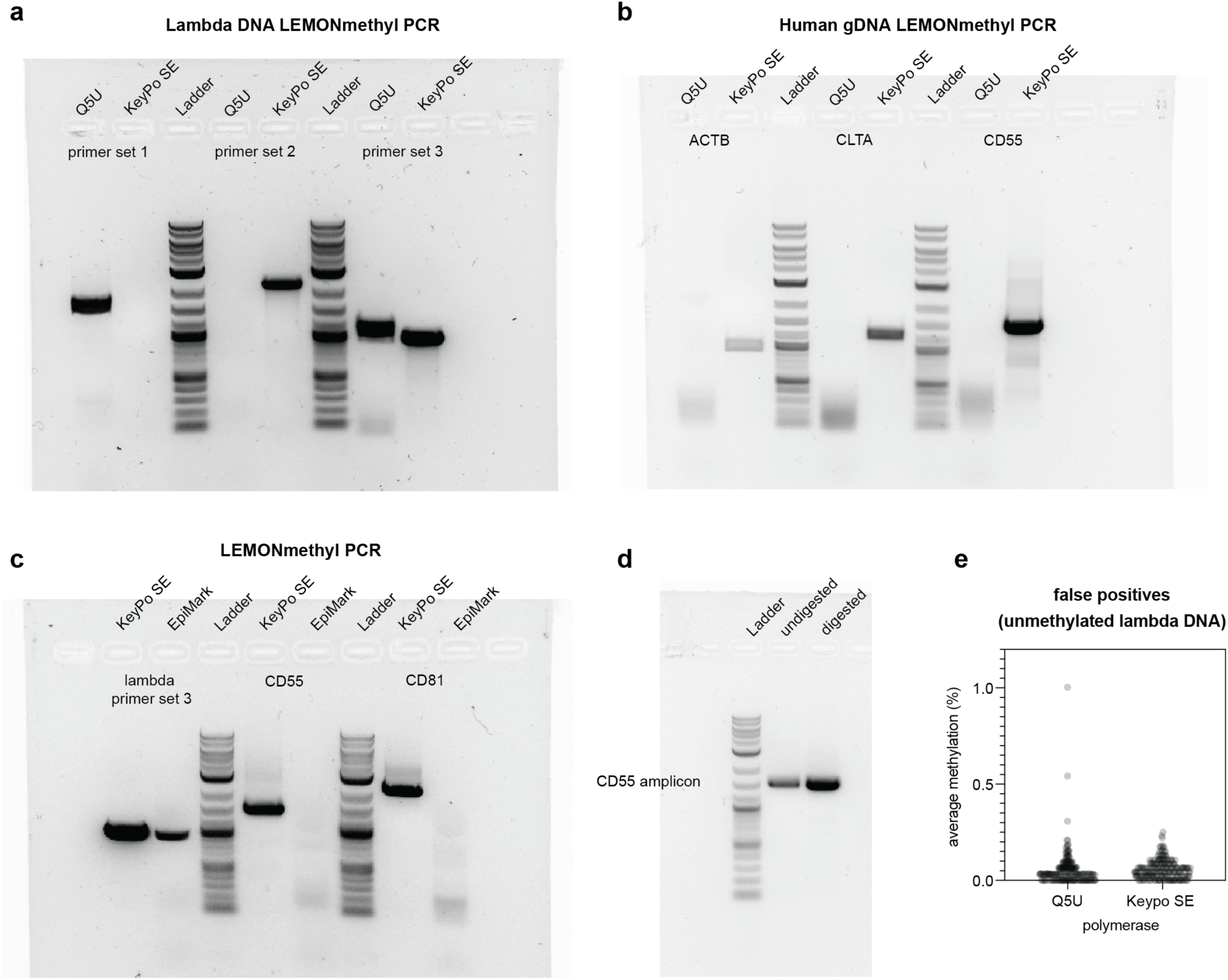
Optimization of LEMONmethyl-seq for robust, long-read DNA methylation profiling. (a) Comparison of Q5U and KeyPo SE polymerases for LEMONmethyl-seq. Agarose gel of LEMONmethyl PCR products generated from three different primer sets amplifying EM-converted unmethylated lambda DNA. (b) Comparison of Q5U and KeyPo SE across three different primer sets amplifying human genomic loci. Agarose gel of PCR products from LEMONmethyl PCR using *ACTB*, *CLTA*, and *CD55* primer sets on EM-converted human genomic DNA. (c) EpiMark Taq polymerase and KeyPo SE comparison. Agarose gel of PCR products from amplification across lambda DNA primer set and two human genomic loci, *CD55* and *CD81*. (d) *CD55* PCR amplification efficiency comparison between undigested and restriction enzyme digested gDNA to enrich the target region before EM conversion. For the digested sample, gDNA was digested with EcoNI and BstXI in buffer r3.1 and purified prior to EM conversion. (e) False positive comparison between LEMONmethyl-seq performed with Q5U and KeyPo SE polymerases. LEMONmethyl-seq workflow was performed on control, unmethylated lambda DNA using lambda primer set #3. Any methylation calling corresponds to a false positive.

**Figure S2.**
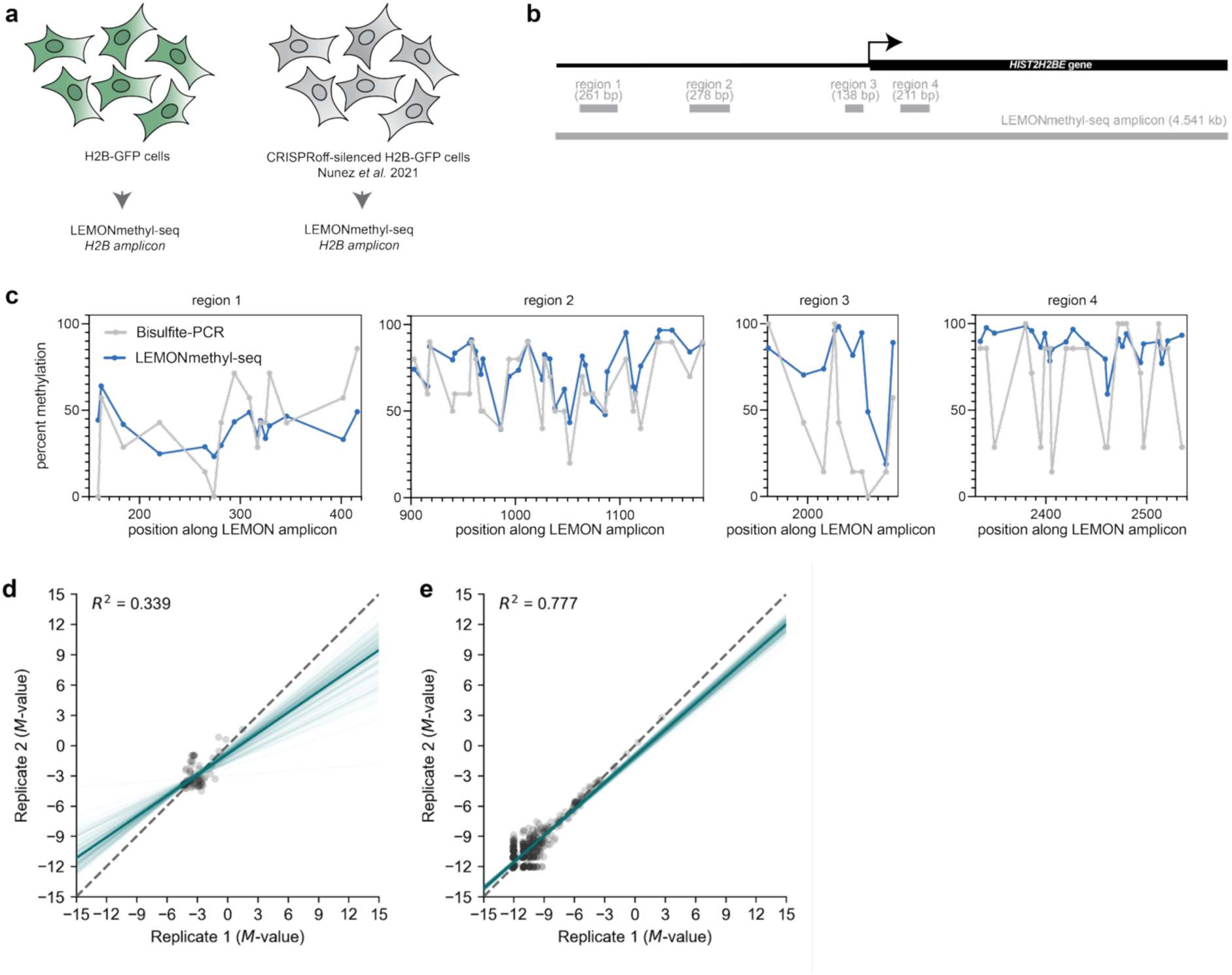
Benchmarking LEMONmethyl-seq to bisulfite-PCR and WGEM-seq. (a) Experimental schematic of LEMONmethyl-seq and bisulfite-PCR benchmarking. H2B-GFP cells from a previous study (Nunez et al. 2021) were harvested for DNA methylation profiling through LEMONmethyl-seq. Both untreated cells and cells with H2B-GFP silenced by CRISPRoff were harvested. (b) Schematic of amplified *H2B* locus. LEMONmethyl-seq amplicon is ∼4.5 kb. Previously, bisulfite-PCR was performed on four different regions, all under 300 bp. (c) Bisulfite-PCR and LEMONmethyl-seq comparison between CRISPRoff-silenced cells. Gray lines represent previous bisulfite-PCR data and blue lines are from LEMONmethyl-seq data. (d) Correlation of *M*-values of each CpG site from *CD55* amplicon between each untreated biological replicate profiled by WGEM-seq. (e) Correlation of *M*-values of each CpG site from *CD55* amplicon between each untreated biological replicate profiled by LEMONmethyl-seq.

**Figure S3.**
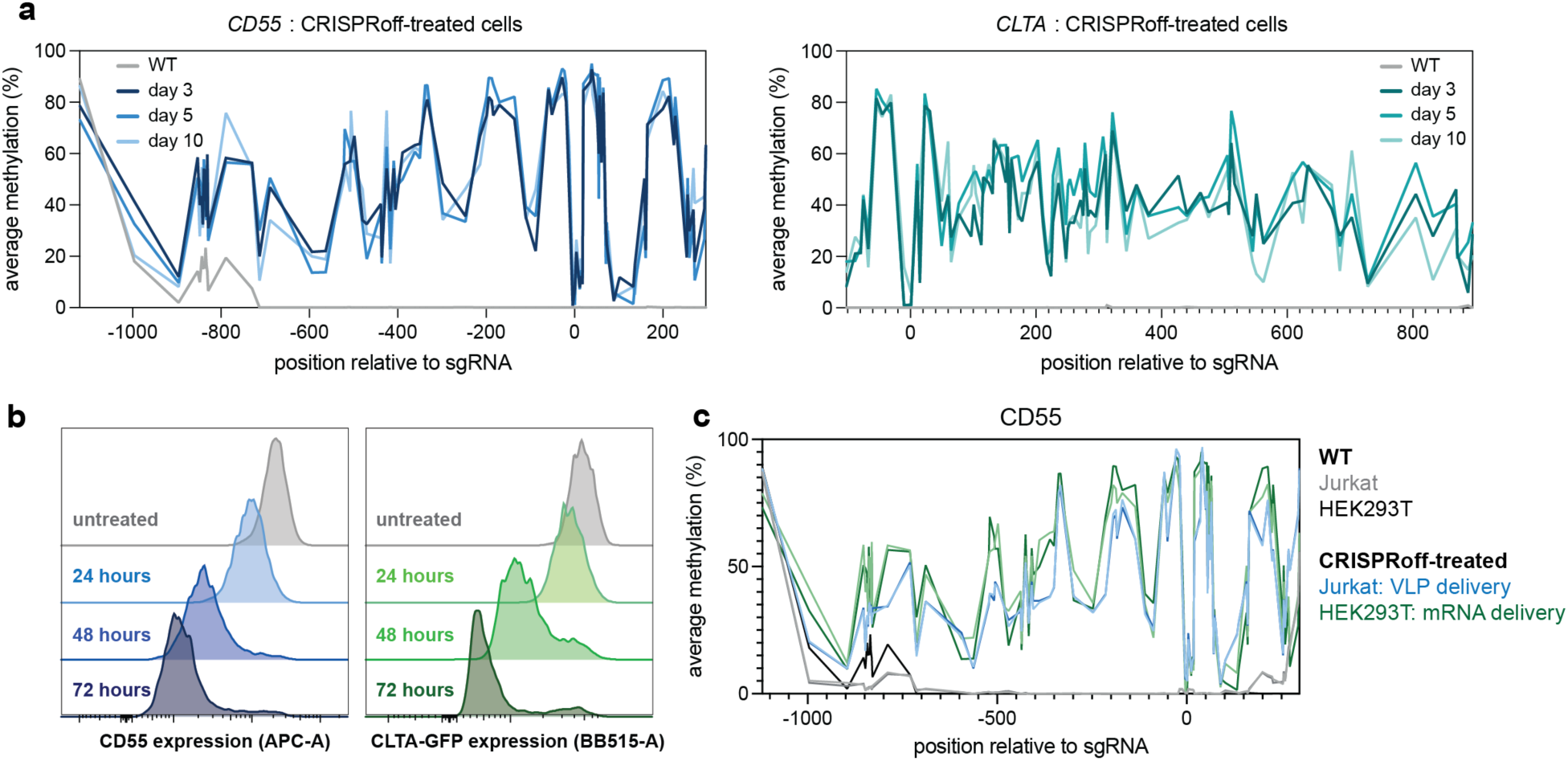
CRISPRoff epigenome editing at *CD55* and *CLTA*. (a) LEMONmethyl-seq profiles of *CD55* and *CLTA* at 3, 5, and 10 days after CRISPRoff mRNA delivery. (b) Flow cytometry histograms of CD55 and CLTA silencing at 24, 48, and 72 hours after CRISPRoff mRNA delivery as described in Fig. 3e. (c) LEMONmethyl-seq data across *CD55* locus in different cell lines and delivery modalities. These data plot *CD55* DNA methylation profiling in Jurkat and HEK293T cells following CRISPRoff-treatment. Methylation profiling in Jurkats is from Fig. 2d (n=2). HEK293T data is from day 3 and 5 following CRISPRoff mRNA treatment from Fig. S3a (n=2).

**Figure S4.**
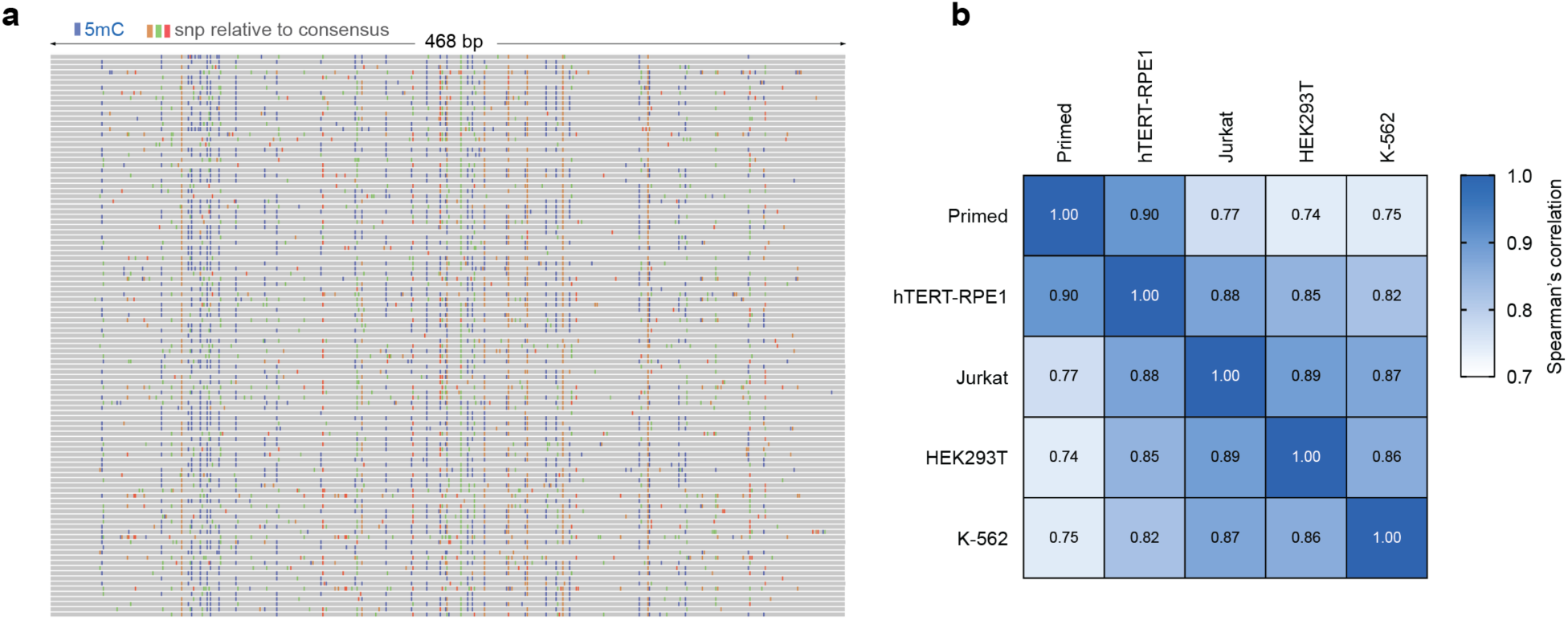
L1Hs 5’ UTR DNA methylation profiling across human cell lines. (a) IGV plot of 100 individual DNA molecules following LEMONmethyl-seq of the consensus L1Hs 5’ UTR amplicon in hTERT-RPE1 cells. 5mC is represented in blue and additional single nucleotide polymorphisms (SNPs) are shown in orange (G), green (A), and red (T), depicting sequence heterogeneity across the L1Hs amplicon. (b) Correlation matrix between L1Hs DNA methylation profiles. Spearman r was calculated comparing the average methylation across the L1Hs amplicon of each cell line.

**Figure S5.**
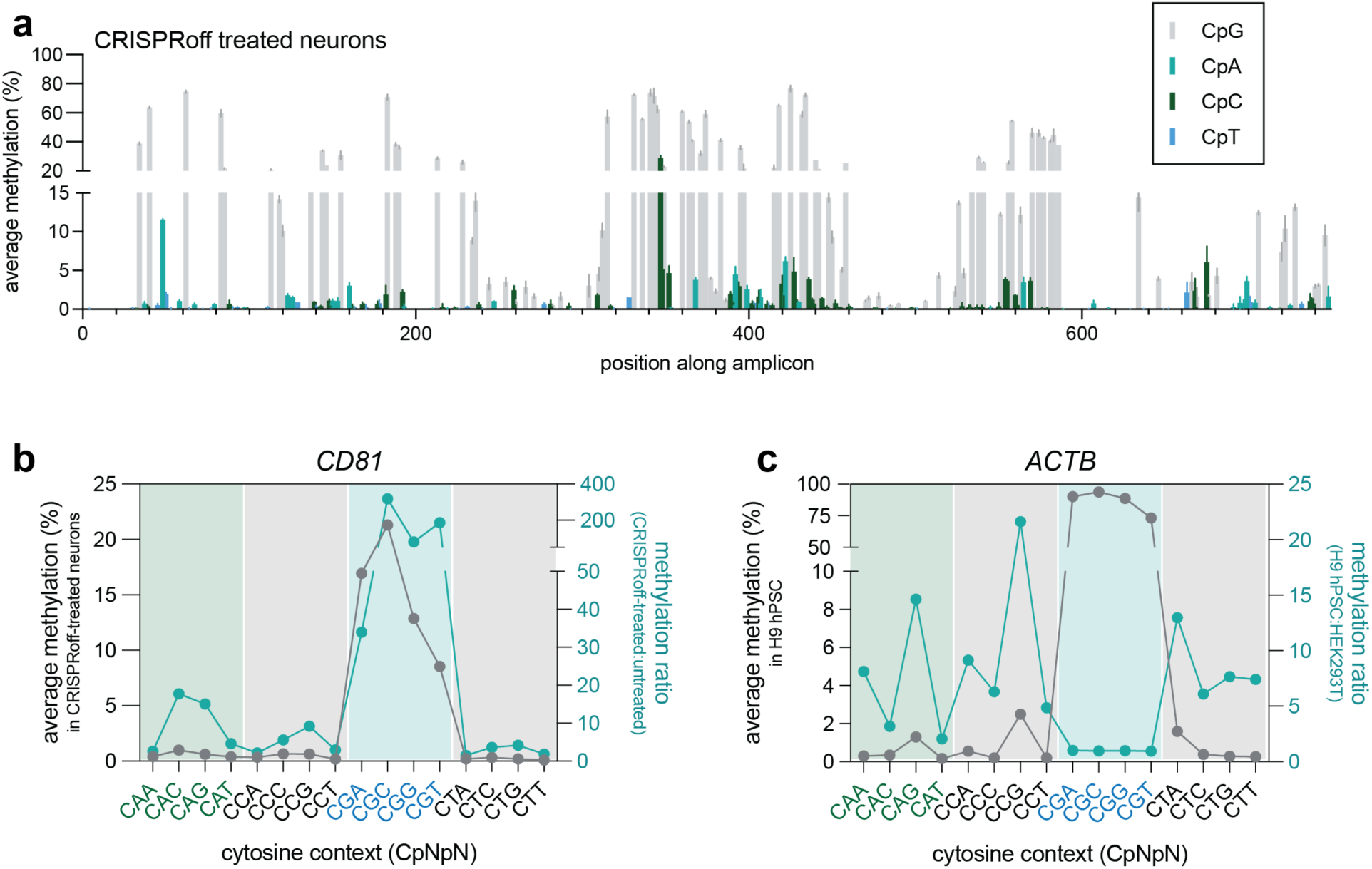
Detection of CpH methylation in neurons and hPSC cells. (a) Bar plot representation of average cytosine methylation across *CD81* amplicon following CRISPRoff treatment in neurons. CpG methylation is shown in gray, CpA methylation in teal, CpC methylation in green, and CpT methylation in blue. Error bars represent standard deviation between two biological replicates. (b) Average methylation and methylation ratio (fold increase in methylation following CRISPRoff treatment relative to untreated neurons) across cytosine contexts. (c) Average methylation and methylation ratio (fold increase in methylation of hPSC *ACTB* compared to HEK293T *ACTB*) across cytosine contexts.

**Table S1. LEMONmethyl-seq oligos.**

Table listing primers used for amplification of EM-converted DNA, amplicon size, optional restriction enzymes for gDNA digestion, and reference sequence. Additionally, each experiment is annotated to indicate whether digested or undigested gDNA was used.

**Table S2. sgRNA and RT-qPCR oligos.**

Table of oligos used for cloning sgRNA protospacers into lentiviral backbones and primers used for RT-qPCR.

**Table S3. Bisulfite-PCR and LEMONmethyl-seq cost comparison.**

Estimates for cost of performing bisulfite-PCR with 10 Sanger reads and cost of performing LEMONmethyl-seq.

**Table S4. PXGGA media composition.**

Description of PXGGA reagent components.

**Table S5. TE consensus sequences from RepBase.**

List of full TE consensus sequences used for primer design.

